# Cohort-specific Boolean models highlight different regulatory modules during Parkinson’s disease progression

**DOI:** 10.1101/2024.02.20.581152

**Authors:** Ahmed Abdelmonem Hemedan, Venkata Satagopam, Reinhard Schneider, Marek Ostaszewski

**Affiliations:** Bioinformatics Core Unit, Luxembourg Centre for Systems Biomedicine, University of Luxembourg, Esch-sur-Alzette, Luxembourg

**Keywords:** Boolean modelling, molecular mechanisms, Disease stratification, Parkinsion’s disease

## Abstract

Parkinson’s Disease (PD) is a multifaceted neurodegenerative disorder characterised by complex molecular dysregulations and diverse comorbidities. It is critical to decode the molecular pathophysiology of PD, which involves complex molecular interactions and their changes over time. Systems medicine approaches can help with this by a) encoding knowledge about the mechanisms into computational models b) simulating these models using patient-specific omics data. This study employs the PD map, a detailed repository of PD-related molecular interactions, as a comprehensive knowledge resource. We aim to dissect and understand the intricate molecular pathways implicated in PD by using logical modelling. This approach is essential for capturing the dynamic interplay of molecular components that contribute to the disease. We incorporate cohort-level and real-world patient data to ensure our models accurately reflect PD’s subtype-specific pathway deregulations. This integration is crucial for addressing the heterogeneity observed in PD manifestations and responses to treatment. To combine logical modelling with empirical data, we rely on Probabilistic Boolean Networks (PBNs).These networks provide a robust framework, capturing the stochastic nature of molecular interactions and offering insights into the variable progression of the disease. By combining logical modelling with empirical data through PBNs, we achieve a more refined and realistic representation of PD’s molecular landscape. The findings provide insights into the molecular mechanisms of PD. We identify key regulatory biomolecules and pathways that differ significantly across PD subtypes. These discoveries have substantial implications for the development of precise medical treatments. The study provides hypothesis for targeted therapeutic interventions by linking molecular dysregulation patterns to clinical phenotypes and advancing our understanding of PD progression and patient stratification.

## 2 Introduction

Parkinson’s disease (PD) is a complex disorder characterized by the progressive degeneration of dopaminergic neurons. This degeneration leads to a variety of motor and cognitive impairments [1]. PD is often accompanied by various comorbidities, such as dementia and diabetes [2], Which further complicate its clinical symptoms and management. An in-depth understanding of PD complexity necessitates a systematic analysis and interpretation of its various subtypes and associated conditions [3].

An important aspect of PD is its molecular pathophysiology. Neuronal degeneration in PD, particularly in the substantia nigra, is closely linked to disruptions in dopamine release, affecting motor cortex stimulation. This molecular perspective is crucial for understanding the disease’s progression and symptoms, such as muscular rigidity, tremor, and bradykinesia. Moreover, the role of molecular mechanisms in the pathogenesis and progression of PD, especially in the context of its comorbidities, remains an important area of exploration.

To investigate the complex interplay of molecular mechanisms in PD, systems biomedicine approaches can be employed [4–6]. These approaches allow for a comprehensive analysis of the disease at a molecular level, integrating various biological data and computational methods. By examining the molecular pathways and their interactions, systems biomedicine provides a holistic view of the disease’s underlying mechanisms [7]. These approaches enable a dynamic analysis of complex molecular networks, crucial for understanding PD progression. This necessitates encoding detailed knowledge and relies on a robust simulation framework for accuracy.

In our approach, we utilize the PD map [8] as a primary knowledge source, and logical modelling as a computational paradigm [9]. The PD map encapsulates extensive knowledge about PD-related mechanisms, offering a crucial tool for visualizing and understanding molecular interactions implicated in the disease. Logical modelling complements this by providing both qualitative and quantitative analyses of disease mechanisms, enabling a deeper understanding of the complex biological systems involved in PD. In our approach, we use the PD map as a knowledge source of PD mechanisms [8]. The PD map represents the largest repository of PD pathways available to date, including detailed knowledge about PD-related mechanisms. It serves as an essential tool for visualising molecular interactions implicated in the disease. The pathway diagrams in the PD map are static, but can be modelled and simulated to understand the dynamics of the represented mechanisms [10].

To accurately model the subtype-specific pathway deregulation in PD, we incorporate cohort and real-world omics data.This integration allows us to examine the heterogeneity of the disease and the specific molecular responses to various perturbations. By integrating this data, our models become more representative of the disease mechanisms in different patient groups. To combine logical modelling with empirical data, we rely on Probabilistic Boolean Networks (PBNs) [9]. PBNs enable us to simulate the impact of molecular dysregulation, thus providing more understanding of the disease pathways. This approach is particularly effective in exploring the complexities of PD pathways and their variations across different disease subtypes [11, 12].

Our analysis identified differential expressed miRNAs, ensuring they were validated and had established interactions in brain tissue. Significant miRNAs showed downregulation in PD and were involved in mitochondrial dysfunction. The enrichment analysis using the PD map highlighted key pathways potentially involved in disease progression. These pathways were converted into dynamic probabilistic Boolean models. Building upon the miRNA data, these models were parameterized to represent the disease cohorts. These models revealed distinct behaviors in key molecular pathways across PD subtypes, including dopamine transcription, PI3K/AKT signaling, FOXO3 activity, mTOR-MAPK signaling, and PRKN mitophagy. There was significant dysregulation in mitochondrial biogenesis and neuron survival in the Parkinsonism group, which was exacerbated by Type 2 diabetes mellitus (T2DM). Additionally, our study revealed notable variations in insulin resistance patterns, with the Prodromal group displaying distinctly different profiles compared to other PD subtypes. differences in autophagy and mitophagy activities were observed, suggesting unique disease mechanisms within each PD subtype. This understanding of PD subtypes and the impact of T2DM comorbidity can help to develop targeted and personalized treatment approaches.

This article is structured as follows: in the next Section we provide an outline of the fundamental aspects of PD, its molecular mechanisms, and the methodologies employed in this study. Next Sections presents methods and results of our study of cohort-specific dynamics in key PD pathways. Finally, we discuss our results and conclude with reflections on the potential impact of our research and directions for future studies in this field.

## 3 Background

Parkinson’s disease (PD) is a complex, chronic, and age-related disorder. PD is characterized by accumulation of alpha-synuclein proteins, leading to the formation of Lewy bodies, which are central to neuronal degeneration [13].This is complicated by mitochondrial dysfunctions, which disrupt cellular energy production, and oxidative stress that damages cellular structures [14]. Further, neuroinflammation contributes to the progressive nature of disease by increasing neural loss. These mechanisms are interconnected and affect each other, which increases the disease complexity [15].

The complexity of PD is not limited to its underlying mechanisms. The disease’s interaction with comorbid conditions, such as Type 2 diabetes mellitus, introduce additional layer of complexity. Studies show that T2DM can exacerbate mitochondrial dysfunction in PD, leading to an accumulation of metabolic byproducts that leads to neuronal death [16]. Further, the use of antidiabetic drugs has shown promise as neuro-protective agents in PD, suggesting common therapeutic pathways for these comorbid conditions [17]. Moreover, studies highlight the interplay between -synuclein pathology, a hallmark of PD, and metabolic dysfunctions characteristic of T2DM, linking these conditions at a molecular level [18]. The complex relationship between PD and its comorbidity highlights the need for a comprehensive cohort data to investigate the diverse and intricate subtypes of this multifaceted disease.

The diversity of PD is reflected in its various subtypes, each with unique molecular signatures. Analysing cohort-level data, such as from the Parkinson’s Progression Markers Initiative (PPMI) cohort study, can help to study the complexities of these subtypes [19]. This approach enables the identification of specific biomarkers in each subtype, allowing for developing precise treatment in PD. The PPMI study provides a rich dataset for analysing PD subtypes, focusing on prodromal, SWEDD (Scans Without Evidence of Dopaminergic Deficit), and parkinsonism cohorts. The prodromal stage represents early PD signs before clear motor symptoms appear. SWEDD patients exhibit PD-like symptoms but lack dopaminergic deficits in scans, suggesting different disease mechanisms. Parkinsonism includes typical PD motor symptoms. In these cohorts, specific miRNAs were identified as potential biomarkers, reflecting the molecular changes associated with each subtype [19].

MicroRNAs (miRNAs) are emerging as significant biomarkers in PD due to their stability in human fluids, and their role in gene regulation [20, 21]. However, this area of research faces challenges due to the non-specific nature of some miRNAs, which are also implicated in other diseases. This limits their specificity for PD diagnosis. Further, the expression levels of key miRNAs can vary based on factors like drug interactions, necessitating careful consideration in experimental designs [22]. These challenges highlight the need for robust and reliable resources like the PD map, which provides a comprehensive and expert-reviewed knowledge base resource.

The PD map serves as a valuable knowledge repository, providing high-quality, disease-specific information crucial for computational modelling. The PD map can be translated into a dynamic Boolean model, allowing for dynamic simulation and predictions. Boolean models offer a simple approach to represent complex disease mechanisms. The biomolecules in a disease mechanism can be represented as model components and their interactions are described by Boolean functions. Boolean models are advantageous as they do not require detailed kinetic information, making them suitable for large-scale data analysis and hypothesis testing. PBNs extend the basic Boolean model by incorporating stochastic elements, which more accurately reflect the inherent variability and dynamic nature of biological systems [9, 23]. In this context, we used pyMaBoss, in which a biological system is depicted as a network of interconnected Boolean variables, each of which represents the state of a biomolecule (e.g., present or absent, active or inactive). Boolean rules, which determine how one variable’s state can influence another’s, define the interactions between these variables. pyMaBoss employed Monte Carlo algorithm operates by randomly sampling possible system states at each time step. This sampling is based on the probabilities of each state given the system’s current state and the rules dictating variable interactions. Through multiple time step simulations, PyMaBoSS can estimate the probability of each state at every point in time to simulate the dynamics of a model [24]. To this end, Boolean modelling has significant applications in clinical and translitional medical research for a range of purposes. Simulations of complex biological systems have enabled prediction of pathway endpoint activities, drug targets, and cellular crosstalks. Identifying attractors has provided an understanding of phenotype activities as they represent steady states of components. Furthermore, attractor comparisons before and after perturbations can shed light on how in vivo systems maintain their homeostasis [25–32].

## 4 Methods

To study the mechanisms of PD, we applied a methodology based on high-quality data and specialized knowledge resources see Figure 1). Using the PPMI dataset and the PD map repository, we employed probabilistic Boolean Modeling techniques. Key genes targeted by the miRNA identified in the PPMI dataset were used to select relevant pathways in the PD map. Then, these pathways were translated into Boolean models (BMs) and parameterized using the cohort data to represent different PD subtypes. Simulation and analysis of these models provided insight into i) differences in pathway dynamics between studied PD subtypes and ii) potential impact of T2DM comorbidity on these pathways

**Fig. 1.**
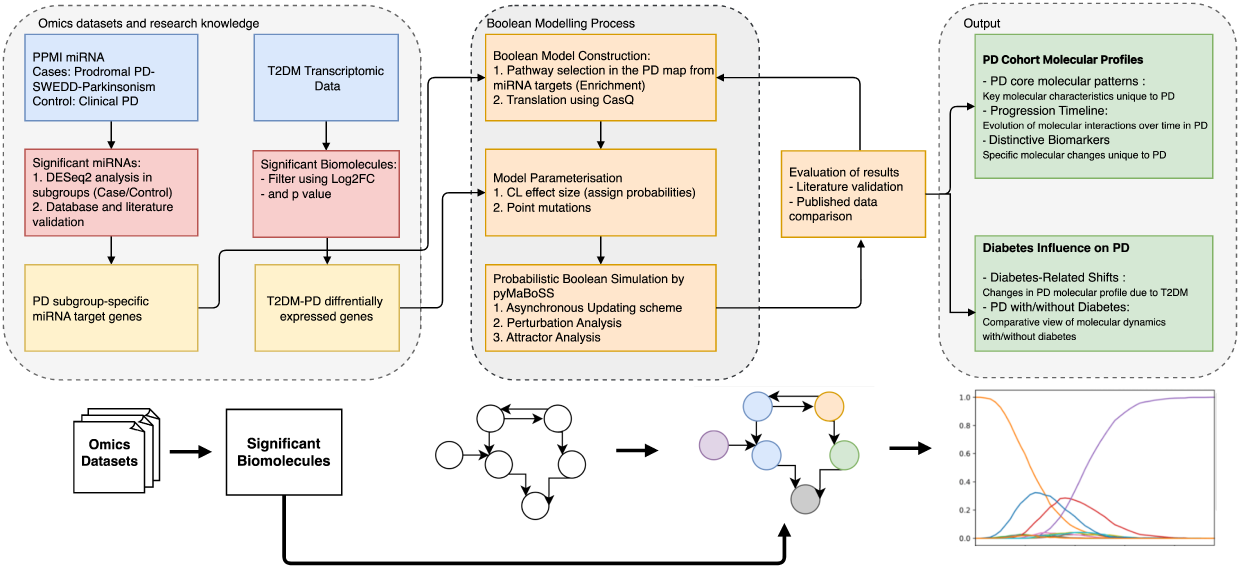
This diagram outlines the research framework for understanding the molecular interactions in Parkinson’s disease groups. We integrate omics datasets and research knowledge to identify significant biomolecules, constructs a Boolean model for simulation, and evaluates the results to determine PD cohort molecular profiles and the influence of diabetes on PD progression.

### 4.1 Parkinson’s Progression Markers Initiative-miRNAs dataset

The Parkinson’s Progression Markers Initiative (PPMI) dataset constitutes a longitudinal observational study of multiple disease cohorts. From the PPMI dataset we used microRNAs derived from blood samples of individuals from cohorts as listed below.

- **Clinical PD:** Describes individuals who exhibit clinical manifestations of PD and possess a positive dopamine transporter (DAT) SPECT scan.
- **Prodromal PD:** Denotes individuals whose symptoms have not yet severely manifested, yet their DAT SPECT results are significantly positive, a group that includes 223 individuals in this dataset.
- **SWEDD:** Describes individuals who, despite a clinical diagnosis of PD, do not exhibit a dopaminergic deficit on their DAT SPECT scan, encompassing 187 individuals in this dataset.
- **Atypical Parkinson’s Disease/Parkinsonism:** Characterizes individuals who exhibit idiopathic symptoms similar to those observed in standard PD.

This dataset is accessible through the Laboratory of Neuro Imaging (LONI) archive at www.ppmi-info.org/data.

### 4.2 Analysis of PPMI dataset

We analysed miRNA expression data from the PPMI dataset, focusing on differentially expressed miRNAs between case and control cohorts. The analysis was conducted using R, a statistical programming language, with specific emphasis on the DESeq2 package for processing and analysing count-based NGS (next-generation sequencing) data. The methodology was structured as follows:

The DESeq2 package in R was used to calculate the log2 fold change (log2FC) in miRNA expression between the case and control groups. We calculated normalised expression values and log2FC values to identify miRNAs with significant changes in expression levels between the case and control cohorts [33]. Further, P value was calculated between the the case and control cohorts using the paired sample t-test [34].

We then identified validated miRNA targets utilizing manually curated databases, with miRTarBase serving as a prominent resource [35]. miRTarBase encompasses over three hundred and sixty thousand experimentally validated miRNA-target interactions. In addition to the miRTarBase, we considered other databases to identify the consensus of miRNA target interactions. These datasets included DIANA-TarBase, miRanda, PicTar and TargetScan which include experimentally validated data with high quality annotation. The overlapped miRNA target interactions were selected for our analysis.

Identified miRNA targets were filtered to match the expression profile of PD substantia nigra. To this end, we only retained miRNAs having at least one target among the SN-expression profile. This profile contains differentially expressed genes from a meta-analysis of eight transcriptomic profiles of post mortem substantia nigra of PD patients vs controls [36], which appear at least once in the PD map.

To further characterize these miRNAs, we compared their expression patterns with curated miRNA expression databases: MiREDiBase [37], miRGate [38], the Human miRNA and Disease Database (HMDD) [39], and through GEO data screening [40].

### 4.3 Calculation of the transcriptomic profile from an organoid model of type Two Diabetes Mellitus

To examine the effect of T2DM on the progression of PD subtypes identified in the PPMI study, we used two transcriptomic datasets that elaborate on PINK1 and GBA mutations in T2DM [41].

In the first dataset, samples from three distinct PD patients carrying a homozygous Q456X mutation in the PINK1 gene were analyzed in comparison to their respective isogenic gene-corrected controls. The dataset comprises differentially expressed genes (DEGs) identified through RNA-sequencing from iPSC-derived neurons after 30 days of differentiation. PINK1 mutant neurons show reduction of IRS1 levels and impaired insulin signaling, indicated by decreased phosphorylation of AKT at S473 and T308. The second dataset elucidates DEGs of the GBA N307S mutation, derived from isogenic control midbrain organoids. The GBA N307S mutation in midbrain organoids is primarily associated with impaired neuron differentiation and cell cycle defects. This mutation shows a significant alteration in lipid metabolism and insulin signalling. We validated the significance of these datasets for the study of PD-T2DM comorbidity as follows:

## 1. Enrichment Analysis

We performed Gene Set Enrichment Analysis (GSEA), utilizing the MINERVA GSEA Plugin,to identify significant pathways within the Parkinson’s Disease (PD) map. Our selection criteria was based on pathways showing signs of insulin resistance and metabolic dysregulations, key elements in PD pathology. Additionally, we used the MsigDB Hallmark 2020 database to identify the molecular signatures relevant to the datasets. These signatures were chosen based on their significant P-values and the relevance of their names known to be associated PD. Further, we performed pathway enrichment analysis using the EnrichNet tool. This tool highlights significant pathways based on significant P-values. Furthermore, we analyzed the brain tissue specificity using InnateDB. Based on the complete set of DEGs, we calculated the XD score metric with a stringent threshold of 0.5, which represents the degree to which a DEG is expressed in a specific tissue type.

## 2. Literature Review

A literature search was conducted to find validated miRNA targets commonly found in both PD and DM. Articles that reported such findings were reviewed and the relevant information was extracted. The results from this literature search were further filtered by the significant targets identified through the enrichment analysis.

## 3. Comparative Analysis

The filtered common miRNA targets were compared with those identified in the PPMI dataset. The aim was to establish a connection between the two diseases prior to proceeding with simulation.

### 4.4 Constructing Boolean models from systems biology diagrams

We constructed Boolean models based on system biology diagrams from the Parkinson’s disease map hosted on the MINERVA Platform [6]. The MINERVA Platform provides the capacity to export selected segments of the map, and we refer to these parts as diagrams. We implemented a stratification process based on diagrams selected through a pathway enrichment analysis of the PPMI dataset using the Parkinson’s disease map. Once we identified significant pathways, we exported these as diagrams in CellDesigner SBML formats for subsequent modelling.

We then converted these models into SBML-qual, a designated module of the SBML standard specifically crafted to represent qualitative models of biological systems. To translate these diagrams into SBML-qual models, we used CaSQ (CellDe-signer as SBML-qual)[42]. We transformed the diagrams into the Simple Interaction Format (SIF) with CaSQ, to create SBML-qual models that are compatible with tools such as BoolNet.

### 4.5 Probabilistic Boolean Model Simulation

The selected Boolean Models (BMs) were simulated using the pyMaBoSS framework, a Python API designed for the MaBoSS software [43] for probabilistic Boolean modeling and simulation. pyMaBoSS leverages continuous time Markov processes. It utilizes a Monte Carlo algorithm for simulating the system’s evolution over time, based on the initial states of the biomolecules and the governing rules of their interactions. pyMaBoSS employs asynchronous updates in a random walk manner, updating a state of a single biomolecule at each step. Random asynchronous transitions are applied within pyMaBoSS to discover steady states and complex attractors from predefined initial states.

#### 4.5.1 Parametrization using PPMI miRNA data

To parameterise the constructed models, we estimated the size of effects for each microRNA using the Cohen distance [44]. This measure provides a standardized mean difference between two distinct groups, taking into account the standard deviation. To make these findings more comprehensible, we converted the Cohen distance into probability values using the Common Language effect size (CL) method, as detailed in works by Ruscio [45] and McGraw [46]. The translated probabilities derived from the Cohen distance were assigned to the initial states of the corresponding miRNA targets within the SBML qual models. These computed probabilities reflect the conditions of various disease subtypes.

Simulations with random walks across probabilistic Boolean Models (BMs) were performed with pyMaBoSS to determine the likelihood of the outcomes (phenotypes) observed from these models. This allowed to examine how certain molecular alterations can affect the likelihood of different disease phenotypes. To identify significant shifts within these simulations, we used a regression technique for detecting multiple change points, as outlined by Lindelov [47].

#### 4.5.2 Parametrization using T2DM transcriptomic profile

The biomolecules of the model were parameterized based on T2DM transcriptomic expression profiles. The conversion of expression profiles into perturbations was achieved by categorizing changes in gene expression as either knockouts or overexpressions. A knockout perturbation involved setting the expression level of a biomolecule to ”zero” indicating a complete loss of function. An overexpression perturbation was represented by setting the expression level to ”one” indicating an increased activity beyond its normal physiological level. The application of knockouts and overexpressions was derived from the two datasets pertaining to mutations in the PINK1 and GBA1 genes. The knockouts and overexpressions were incorporated into the Boolean models as point mutations to simulate specific genetic alterations. The ”point mutation” term refers to in silico targeted modifications in the biomolecules of the model. These targeted alterations are permanent changes in the state of the model biomolecules.

#### 4.5.3 Comparison of simulation trajectories

To quantify and compare two simulaton trajectories, we used Dynamic Time Warping (DTW) [48]. DTW operates by dividing the time series into points and measuring the distance between corresponding points in different series. In this context, DTW was used to quantify variability of simulation trajectories based on calculated change points. DTW calculates the distance between each point in one series and every point in the other series, identifying the optimal path that minimizes the total distance between the series. A lower DTW score indicates a higher similarity between the series(fig. 3). A lower DTW score suggests greater similarity between series. Thus, the DTW score can be used as a measure of the ”activity” or dynamics of a particular process or trajectory over time. A lower DTW score could suggest a higher level of activity or more consistent condition progression, while a higher DTW score could indicate less activity or more variability in the process progression. However, interpreting the DTW score requires an understanding of the specific characteristics of the disease conditions. Pearson correlation was utilized to measure the correlation between DTW similarity values in pairs of subgroups (**??**).

## 5 Results

To study complex molecular mechanisms of PD, we analysed PPMI cohort data to focus on significant pathways associated with the PD pathogenesis. We used this information to construct and parametrise Boolean models. Once the models were analysed, we compiled, interpreted and validated our findings. The validation process involved the consistency of the model results with the existing literature to ensure their alignment with the known biological behavior and experimental data. We identified pathways specific for mitochondrial dysfunction and insulin resistance, common across various PD subgroups and T2DM comorbidities, highlighting their critical roles in disease pathogenesis.

### 5.1 Analysis of expression profiles for pathway modelling

#### miRNA data analysis for PD subgroups

We calculated the differential expression of miRNAs and their standardized effect sizes, and transformed them into probabilities using the Common Language Effect Size method. Based on manually curated miRNA databases we identified miRNA targets. The targets were filtered based on the substantia nigra dataset [36] and compared with those reported in published literature and datasets. We found that most of the significant miRNAs were downregulated in PD and were involved in mitochondrial dysfunction. The miRNAs that did not match with the published validated experiments were not considered in further analysis (Table 1).

**Table 1.** The table indicate the matched (top) and mismatched (bottom) expressions between the filtered PPMI-miRNAs and the reported miRNAs in literature and datasets. The table includes the miRNA name, the direction of regulation (up or down), the sample type, the method used for measurement, and the reference for the data.

|  | miRNA | Regulation | Sample | Method | Ref. |
| --- | --- | --- | --- | --- | --- |
| Matched expressions | hsa-miR-96-5p | Up | Peripheral blood | RT-qPCR (TaqMan) | [49] |
|  | hsa-miR-26a-5p | Up | Peripheral blood | RT-qPCR (SYBR Green) | [50] |
|  | hsa-miR-424-5p | Up | Peripheral blood | Microarray | [50] |
|  | hsa-miR-9-3p | Up | Peripheral blood | RT-qPCR (TaqMan) | [50] |
|  | hsa-miR-454-3p | Up | Peripheral blood | RT-qPCR (TaqMan) | [51] |
|  | hsa-miR-15b-5p | Up | Peripheral blood | RT-qPCR (TaqMan) | [51] |
|  | hsa-miR-671-5p | Up | Peripheral blood | Microarray | [52] |
|  | hsa-miR-93-5p | Up | Prefrontal cortex | Illumina's HiSeq 2000 | [52] |
|  | hsa-miR-195-5p | Up | Peripheral blood | RT-qPCR (TaqMan) | [53] |
|  | hsa-miR-20a-5p | Up | Peripheral blood | Microarray | [53] |
|  | hsa-miR-16-5p | Up | Prefrontal cortex | Illumina's HiSeq 2000 | [53] |
|  | hsa-miR-132-3p | Up | Peripheral blood | RT-qPCR (TaqMan) | [54] |
|  | hsa-miR-196b-5p | Down | Peripheral blood | RT-qPCR (TaqMan) | [54] |
|  | hsa-miR-92b-3p | Down | Peripheral blood | Microarray | [54] |
|  | hsa-miR-19a-3p | Down | Mid-brain | Microarray | [54] |
|  | hsa-miR-19a-3p | Down | Peripheral blood | Microarray | [54] |
|  | hsa-miR-92a-3p | Down | Peripheral blood | RT-qPCR (TaqMan) | [54] |
|  | hsa-miR-133b | Down | Mid-brain | RT-qPCR (TaqMan) | [54] |
|  | hsa-miR-15b-5p | Down | Peripheral blood | RT-qPCR (TaqMan) | [54] |
|  | hsa-miR-7-5p | Down | Peripheral blood | RT-qPCR (TaqMan) | [49] |
|  | hsa-miR-15a-5p | Down | Peripheral blood | Microarray | [55] |
|  | hsa-miR-19b-3p | Down | Peripheral blood | RT-qPCR (TaqMan) | [56] |
|  | hsa-miR-139-5p | Down | Peripheral blood | RT-qPCR (TaqMan) | [56] |
|  | hsa-miR-450b-5p | Down | Peripheral blood | RT-qPCR (TaqMan) | [56] |
|  | hsa-miR-212-3p | Down | Prefrontal cortex | Illumina's HiSeq 2000 | [56] |
|  | hsa-miR-22-3p | Down | Peripheral blood | RT-qPCR (TaqMan) | [56] |
|  | hsa-miR-26a-5p | Down | Peripheral blood | RT-qPCR (TaqMan) | [56] |
|  | hsa-miR-16-2-3p | Down | Prefrontal cortex | Illumina's HiSeq 2000 | [56] |
|  | hsa-miR-16-2-3p | Down | Peripheral blood | RT-qPCR (TaqMan) | [57] |
|  | hsa-miR-30b-5p | Down | Peripheral blood | RT-qPCR (TaqMan) | [57] |
|  | hsa-miR-144-3p | Down | Prefrontal cortex | Illumina's HiSeq 2000 | [58] |
|  | hsa-miR-323a-3p | Down | Peripheral blood | RT-qPCR (TaqMan) | [59] |
|  | hsa-miR-495-3p | Down | Peripheral blood | RT-qPCR (TaqMan) | [59] |
|  | hsa-miR-148b-3p | Down | Peripheral blood | RT-qPCR (TaqMan) | [59] |
|  | hsa-miR-374a-5p | Down | Peripheral blood | RT-qPCR (TaqMan) | [59] |
|  | hsa-miR-199b-3p | Down | Peripheral blood | RT-qPCR (TaqMan) | [59] |
|  | hsa-miR-374b-3p | Down | Peripheral blood | RT-qPCR (TaqMan) | [59] |
|  | hsa-miR-20a-5p | Down | Peripheral blood | Microarray | [59] |
| Mismatched expressions | hsa-miR-199b-3p | Up | Peripheral blood | Microarray | [59] |
|  | hsa-miR-196b-5p | Up | Peripheral blood | RT-qPCR (TaqMan) | [60] |
|  | hsa-miR-221-3p | Up | Peripheral blood | RT-qPCR (TaqMan) | [60] |
|  | hsa-miR-103a-3p | Up | Peripheral blood | RT-qPCR (TaqMan) | [60] |
|  | hsa-miR-320b | Up | Prefrontal cortex | Illumina's HiSeq 2000 | [60] |
|  | hsa-miR-30c-5p | Down | Peripheral blood | RT-qPCR (TaqMan) | [49, 60] |
|  | hsa-miR-30a-5p | Down | Peripheral blood | RT-qPCR (SYBR Green) | [57, 60] |
|  | hsa-miR-181c-5p | Down | Peripheral blood | RT-qPCR (TaqMan) | [61] |
|  | hsa-miR-338-5p | Down | Prefrontal cortex | Illumina's HiSeq 2000 | [62] |
|  | hsa-miR-148b-3p | Down | Prefrontal cortex | Illumina's HiSeq 2000 | [63] |
|  | hsa-miR-21-5p | Down | Peripheral blood | Microarray | [50, 63] |

The majority of the identified miRNAs are dysregulated in all cohorts (see Table 1). The effect size of the filtered miRNAs differs between cohorts (see Figure 2).

**Fig. 2.**
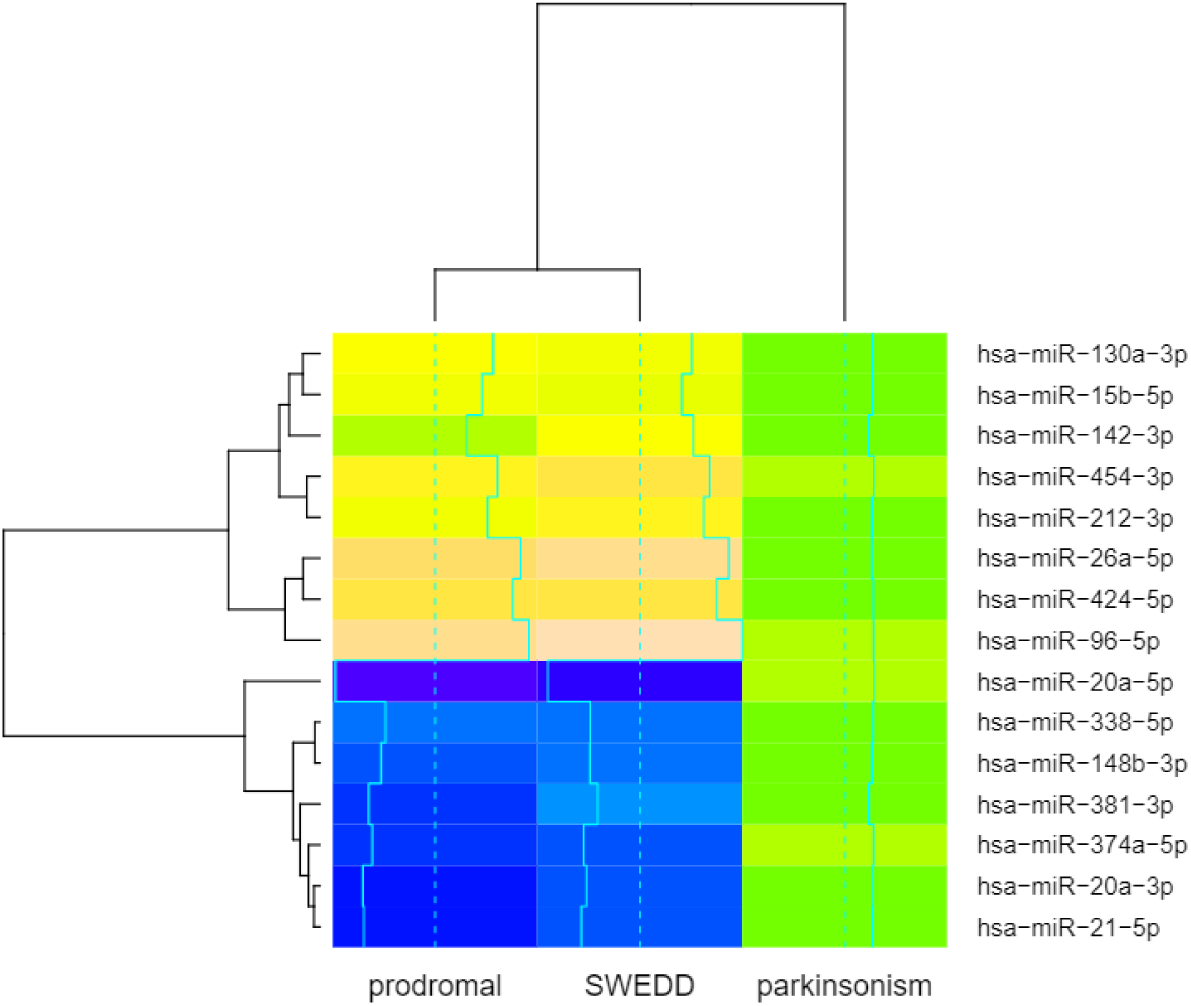
The figure shows an example of significant miRNAs that are common across the conditions with different expressions

#### RNAseq data analysis for T2DM comorbidity

To identify potential connections between PD and T2DM, we analyzed transcriptomic data from PINK1 and GBA mutations in T2DM organoid models and compared them to a substantia nigra dataset of PD, as described in Section 3.2. The following results were obtained:

1. Differentially expressed genes in two datasets describing the PINK1 Q456X and GBA N307S mutations in T2DM were identified and common overlaps with the genes in the substantia nigra dataset on the PD Map were determined [11]. It was found that 81 genes are commonly altered across datasets (see the Suplementary File)
2. Validated common miRNA-target pairs that were reported in the literature as being involved in both PD and T2DM were identified (Table 2) through enrichment analysis of the differentially expressed genes (DEGs) and a literature search. Among the significant DEGs analyzed, a subset of 20 was found to overlap with the substantia nigra dataset.

**Table 2.** The table indicates the common miRNA expression and target regulation in PD and T2DM. The table lists miRNA, targeted genes, and references for studies that have common altered expression of the miRNA in PD or T2DM.

| miRNA | Targets | References |  |
| --- | --- | --- | --- |
|  |  | PD | T2DM |
| hsa-miR-423-3p | CDKN1A | [64] | [65] |
| hsa-miR-132-3p | MAPK1 | [64] | [66, 67] |
| hsa-mir-15a-5p | RET, PHLPP1 | [64] | [68, 69] |
| hsa-mir-29c-3p | PTEN | [70] | [71] |
| hsa-mir-29a-3p | IGF1 | [70, 72] | [71, 73] |
| hsa-mir-20a-5p | PTEN, E2F1 | [64, 74] | [75, 76] |
| hsa-mir-22-3p | PTEN | [77] | [78] |
| hsa-mir-26b-5p | IGF1R, PTEN | [79] | [80] |
| hsa-mir-143-3p | IGF1R, AKT1 | [64] | [81] |
| hsa-mir-145-5p | IGF1R, IRS1, EIF4E, RPS6KB1 | [82] | [83] |
| hsa-mir-133b | IGF1R, AKT1 | [84] | [85] |
| hsa-mir-34a-5p | E2F1 | [86] | [87] |
| hsa-mir-182-5p | PTEN, GSK3B | [88] | [89] |
| hsa-mir-148a-3p | IRS1 | [79] | [90] |
| hsa-mir-7-5p | SNCA, IGF1R, RS1 | [64, 79] | [91] |
| hsa-mir-195-5p | RET, INSR | [79] | [92] |
| hsa-mir-218-5p | RET | [93] | [94] |
| hsa-mir-200c-3p | ROCK2 | [74] | [95] |
| hsa-miR-125b-2-3p | IGF1R | [96] | [97] |
| hsa-mir-18a-5p | PTEN, PHLPP1 | [74] | [98] |
| hsa-miR-17-5p | PHLPP1, PTEN, E2F1 | [93] | [99] |
| hsa-mir-96-5p | GSK3B | [100] | [101] |
| hsa-mir-21-5p | PTEN, E2F1 | [102] | [103] |
| hsa-mir-200a-3p | PTEN, MAPK14 | [104] | [105] |
| hsa-miR-200b-3p | PHLPP1, ROCK2 | [104] | [105] |
| hsa-miR-200c-3p | ROCK2 | [104] | [105] |
| hsa-mir-103a-3p | PTEN | [64] | [105] |
| hsa-mir-10a-5p | PTEN | [21] | [106] |
| hsa-mir-153-3p | SNCA, PTEN | [21] | [107] |
| hsa-mir-19b-3p | PTEN | [21] | [108] |
| hsa-mir-155-3p | PTEN | [72] | [109] |
| hsa-mir-26a-5p | GSK3B, PRKCD, PTEN | [72] | [110] |
| hsa-miR-26b-5p | IGF1R, PTEN | [72] | [110] |
| hsa-let-7a-5p | E2F1 | [72] | [111] |
| hsa-mir-23a-3p | PTEN | [77] | [112] |
| hsa-mir-100-5p | AKT1, IGF1R | [113, 114] | [115] |
| hsa-mir-92a-3p | PHLPP1, PTEN | [113] | [116] |

To identify pathways plausible for Boolean modelling, we performed enrichment analysis for the targets of miRNAs identified in the PPMI cohort. Using the PD map as a pathway repository, this analysis produced significant pathways as listed in Table 3. These pathways were translated into Boolean models (BMs) and subsequently parameterized in accordance with effect size computations (Table 5).

**Table 3.**
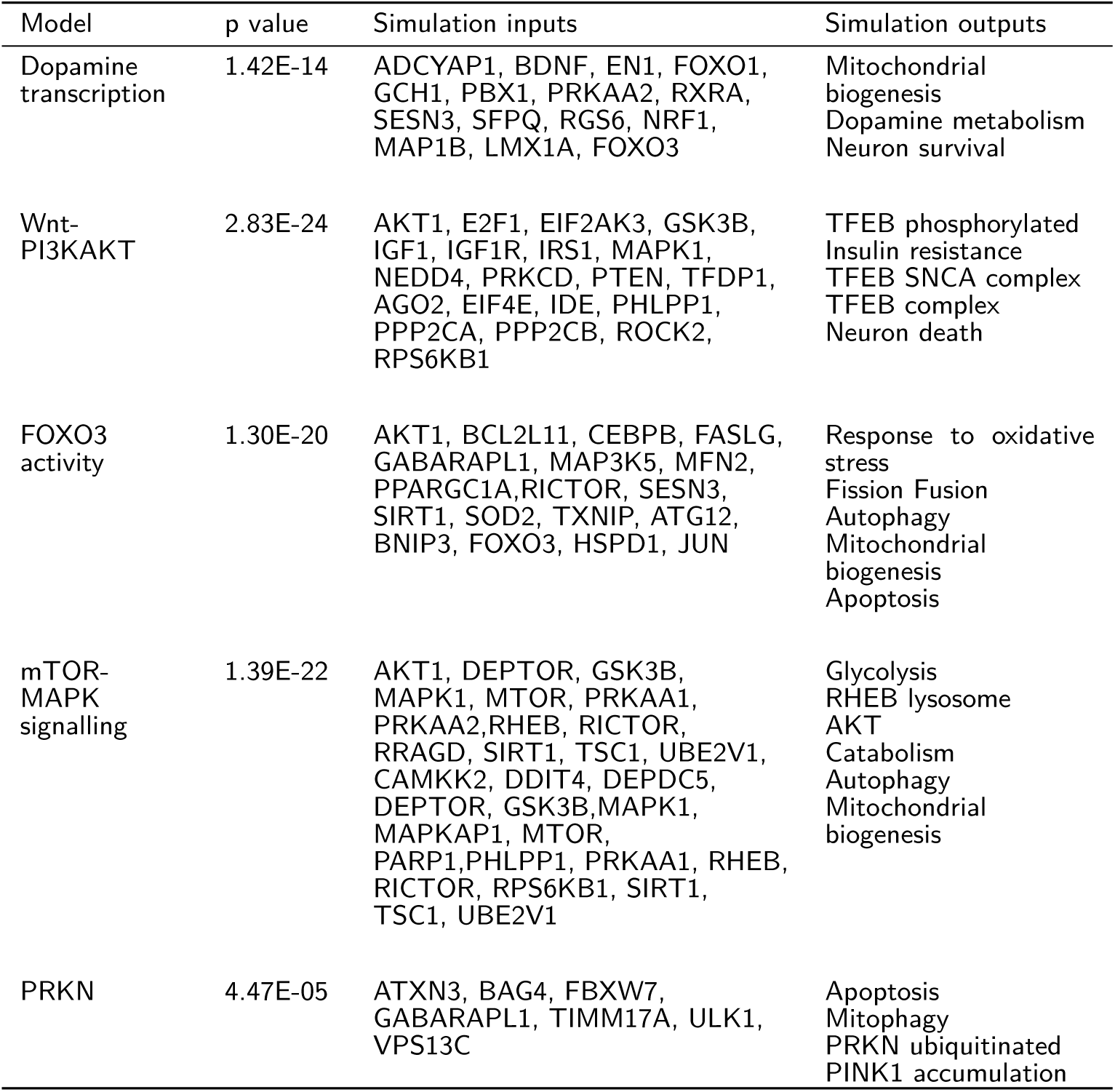
This table presents the Boolean models, with the inputs and outputs for each model listed. The pathways included are the Dopamine transcription pathway, the Wnt-PI3KAKT pathway, the FOXO3 activity pathway, and the mTOR-MAPK signalling pathway. The inputs for each pathway consist of various biomolecules, while the outputs represent various cellular processes or biomolecules that are influenced by the inputs.

| Model | p value | Simulation inputs | Simulation outputs |
| --- | --- | --- | --- |
| Dopamine transcription | 1.42E-14 | ADCYAP1, BDNF, EN1, FOXO1, GCH1, PBX1, PRKAA2, RXRA, SESN3, SFPQ, RGS6, NRF1, MAP1B, LMX1A, FOXO3 | Mitochondrial biogenesis<br>Dopamine metabolism<br>Neuron survival |
| Wnt-PI3KAKT | 2.83E-24 | AKT1, E2F1, EIF2AK3, GSK3B, IGF1, IGF1R, IRS1, MAPK1, NEDD4, PRKCD, PTEN, TFDP1, AGO2, EIF4E, IDE, PHLPP1, PPP2CA, PPP2CB, ROCK2, RPS6KB1 | TFEB phosphorylated<br>Insulin resistance<br>TFEB SNCA complex<br>TFEB complex<br>Neuron death |
| FOXO3 activity | 1.30E-20 | AKT1, BCL2L11, CEBPB, FASLG, GABARAPL1, MAP3K5, MFN2, PPARGC1A, RICTOR, SESN3, SIRT1, SOD2, TXNIP, ATG12, BNIP3, FOXO3, HSPD1, JUN | Response to oxidative stress<br>Fission Fusion<br>Autophagy<br>Mitochondrial biogenesis<br>Apoptosis |
| mTOR-MAPK signalling | 1.39E-22 | AKT1, DEPTOR, GSK3B, MAPK1, MTOR, PRKAA1, PRKAA2, RHEB, RICTOR, RRAGD, SIRT1, TSC1, UBE2V1, CAMKK2, DDIT4, DEPDC5, DEPTOR, GSK3B, MAPK1, MAPKAP1, MTOR, PARP1, PHLPP1, PRKAA1, RHEB, RICTOR, RPS6KB1, SIRT1, TSC1, UBE2V1 | Glycolysis<br>RHEB lysosome<br>AKT<br>Catabolism<br>Autophagy<br>Mitochondrial biogenesis |
| PRKN | 4.47E-05 | ATXN3, BAG4, FBXW7, GABARAPL1, TIMM17A, ULK1, VPS13C | Apoptosis<br>Mitophagy<br>PRKN ubiquitinated<br>PINK1 accumulation |

### 5.2 Construction of models of Parkinson’s disease pathways

We constructed Boolean models based on the PPMI cohort-specific dataset and the PD map. This helped us choose the significant diagrams for further analysis and modelling. The criteria for selecting diagrams for subsequent modeling and stratification were based on the pathway enrichment analysis of the PPMI dataset using the PD map. The enriched pathways included dopamine transcription pathways, PI3k/AKT signalling, FOXO3 activity, mTOR-MAPK signalling, and PRKN mitophagy. These pathways emphasize that the consequences of their dysregulation are largely dependent on the characteristics of the disease subgroups. To ensure comprehensive coverage of all disease subtypes under consideration, we incorporated all miRNA targets from the PPMI dataset into the enrichment analysis.

The pathways identified via the enrichment analysis were exported from the PD map in CellDesigner SBML format and translated into SBML-qual files by CaSQ tool. These SBML-qual files were first verified for correctness and completeness. To this end we performed structural and dynamical analysis. Structural verification involved the assessment of interactions among the biomolecules of the model. The assesment examined the interactions between the biomolecules, focusing on the nature of the interactions. The examination was achieved by using SIGNOR database to verify the type of the interactions-whether inhibitory or stimulatory. Additionally, the assessment involved identification of the directionality of these interactions using the same database.

Dynamic verification examined the model’s dynamic behavior over iteration steps, evaluating the model response against single perturbations. We altered the state of single nodes to observe the effects on the model behaviour. We validated the reliability of our BMs behaviour by comparing their behavior with actual data. Through simulations, we assessed the capacity of BMs to mimic known perturbations, and their reliability in modelling corresponding biological processes. The simulated behavior of these pathways matched the expected behavior according to published literature (See Table 4). The simulated pathways’ responses were compared to expected biological outcomes, and the coherence of dynamic patterns observed in literature (see **??**. We conducted the comparison by examining each model’s simulated responses that reflect the expected biological behaviors. The metrics to decide the the model behaved correctly were qualitative measures. We aligned the ON/OFF state transitions of each biomolecule in the simulation with the corresponding activation/inhibition behavior described in the literature. We considered the model accurate when an ON state corresponded to activation behavior and an OFF state corresponded to inhibition, as per established biological findings..

**Table 4.**
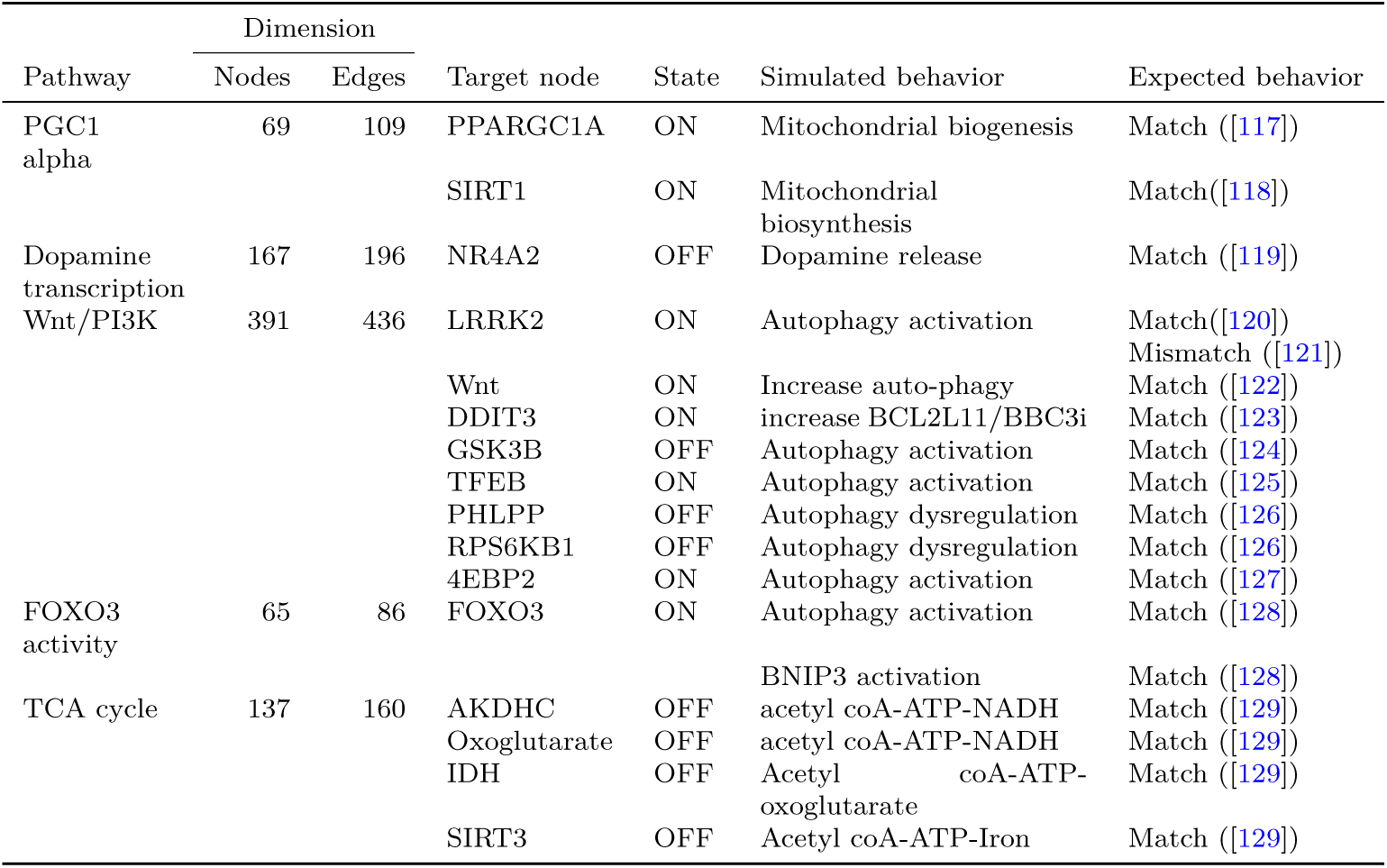
The table compares the simulated behavior of several Boolean models to expected behavior based on published literature. The table includes information on the pathways, the number of nodes and edges in each network, the target node, the state of the target node (ON or OFF), and the simulated and expected behavior for each pathway.

### 5.3 Model parameterisation using cohort data

Using the miRNA expression data of the PPMI dataset we determined the probabilities of the initial states of BMs, based on the effect size and statistical correlations of each subgroup with the control (PD clinical). We used these calculated probabilities in BM models constructed from the PD map to run simulations in pyMaBoSS.

With these simulations of BM we explored the likelihoods associated with reaching levels of specific model outputs, listed in Table 3. With this we evaluated the impact of molecular changes on the states of cellular phenotypes [28, 130, 131].

The illustration in Table 5 shows the computed miRNA effect sizes specific to the SWEDD, prodromal, and parkinsonism subtypes, along with associated targets. When this tailored model is simulated using pyMaBoSS, it yields output readouts for the three phenotypes(see Figure 3).

**Table 5.**
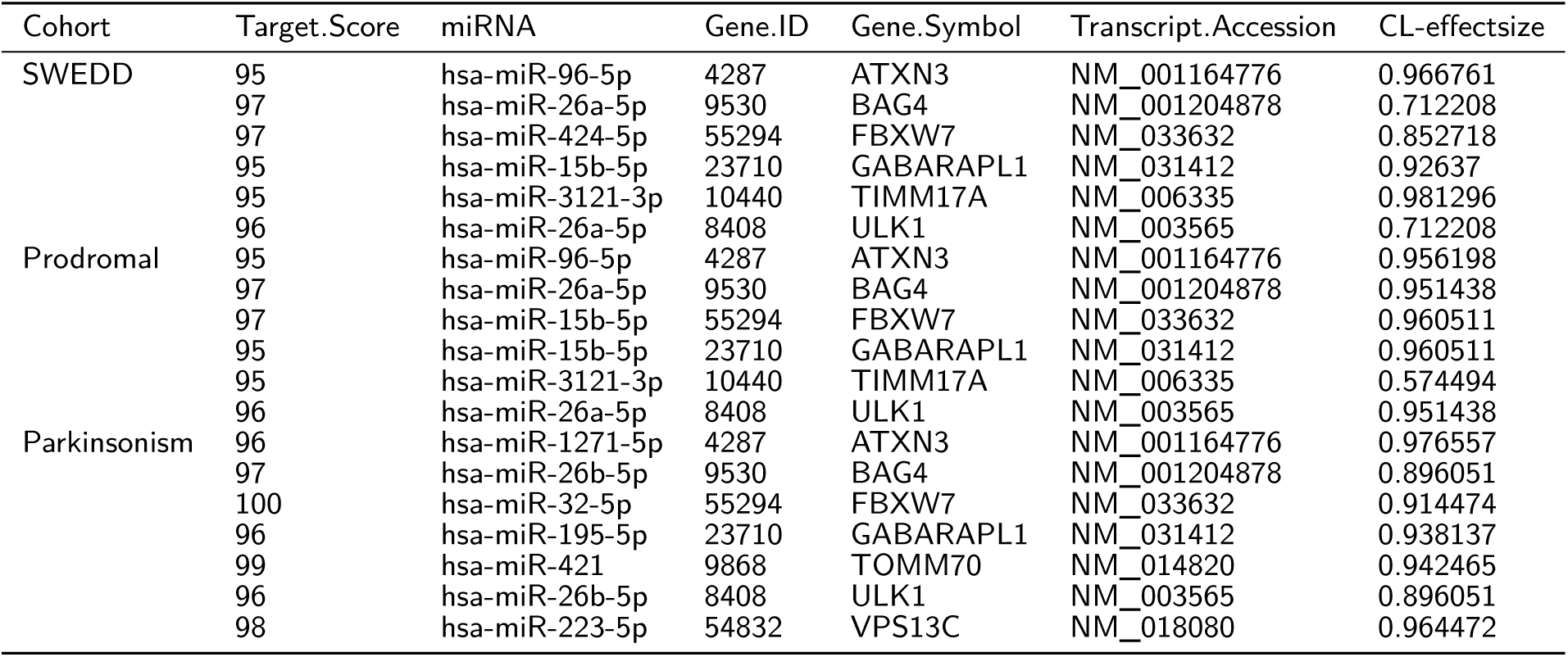
This table presents PRKN-mitophagy Boolean model parameters within three subgroups SWEDD, prodromal, parkinsonism with common language (Cl) effect sizes and targets

**Fig. 3.**
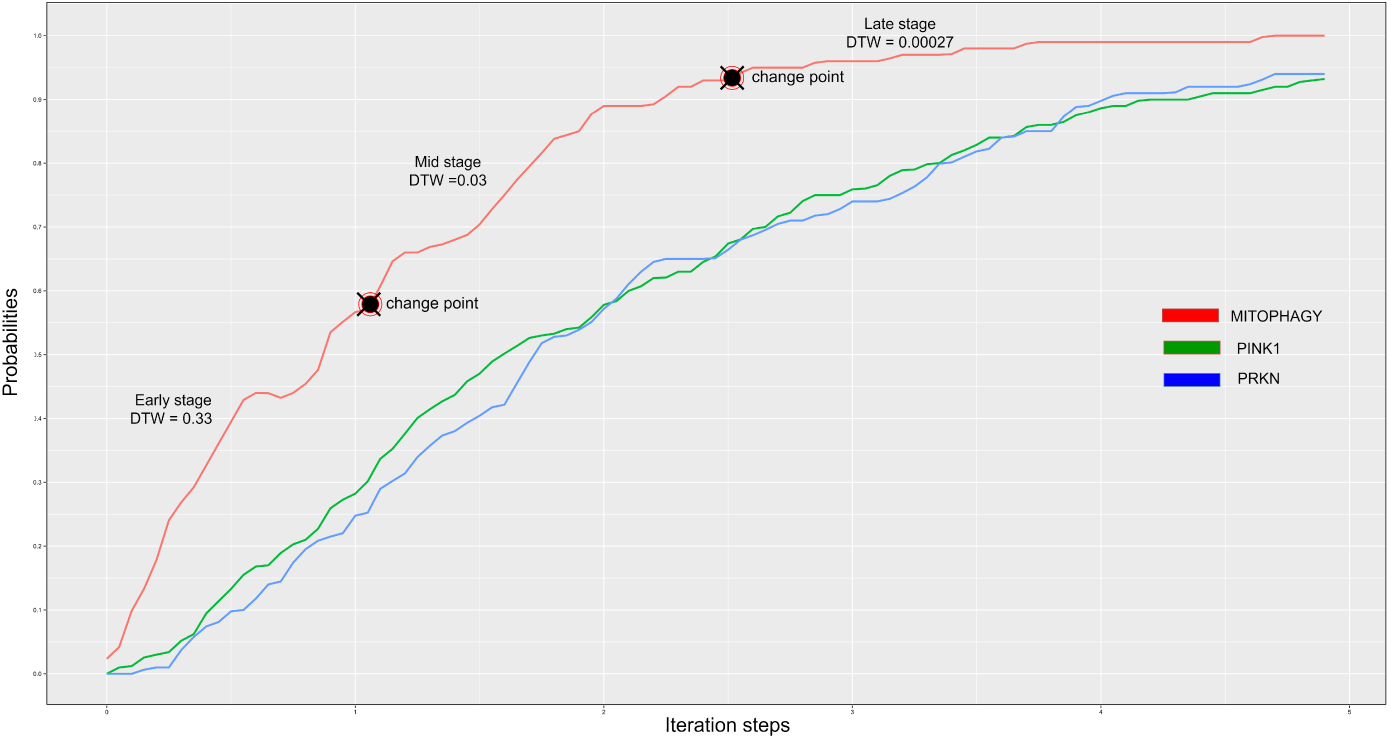
The figure shows a representative run of the simulation in PRKN mitophagy model, consisting of 100 iteration steps with 100 repetitions. Dynamic time warping measures similarity between two sequences which differs in speed based on different stages of the simulation (early, mid, and late).

In the simulated graphs shown in (Figure 3), we calculated their self-similarity and compared them pairwise to other simulates conditions, as described in Methods (Section 3.5.3). Through the process of comparing graphs pairwise, we can identify which aspects of the system remain consistent across different simulations and which ones vary. This helps in pinpointing recurring or unique patterns within the simulation, which are key to understanding the model behavior. For instance, if two simulated endpoints of a biological system reveal similar patterns despite changes in certain parameters. The observed differences between the simulations can highlight the impact of specific parameters on model endpoints in different groups.

The selected models were additionally parameterized to investigate the comorbidity of PD and T2DM. Utilizing the DEGs identified from the T2DM datasets, perturbations were introduced within these models.

DEGs identified with high expression levels were set in the models as permanently activated (simulated overexpression), and downregulated DEGs were set as permanently inactive (simulated knockout). As illustrated in Table 6, the model of PRKN pathway simulating PD-T2DM comorbidity uses two sets of parameters: i) miRNA-based and ii) specific to T2DM. During simulation, these sets of parameters are used together. The miRNA-based parameters represent characteristics of a given cohort, and the T2DM-specific parameters encode the impact of T2DM on the evolution of the PD cohort.

**Table 6.**
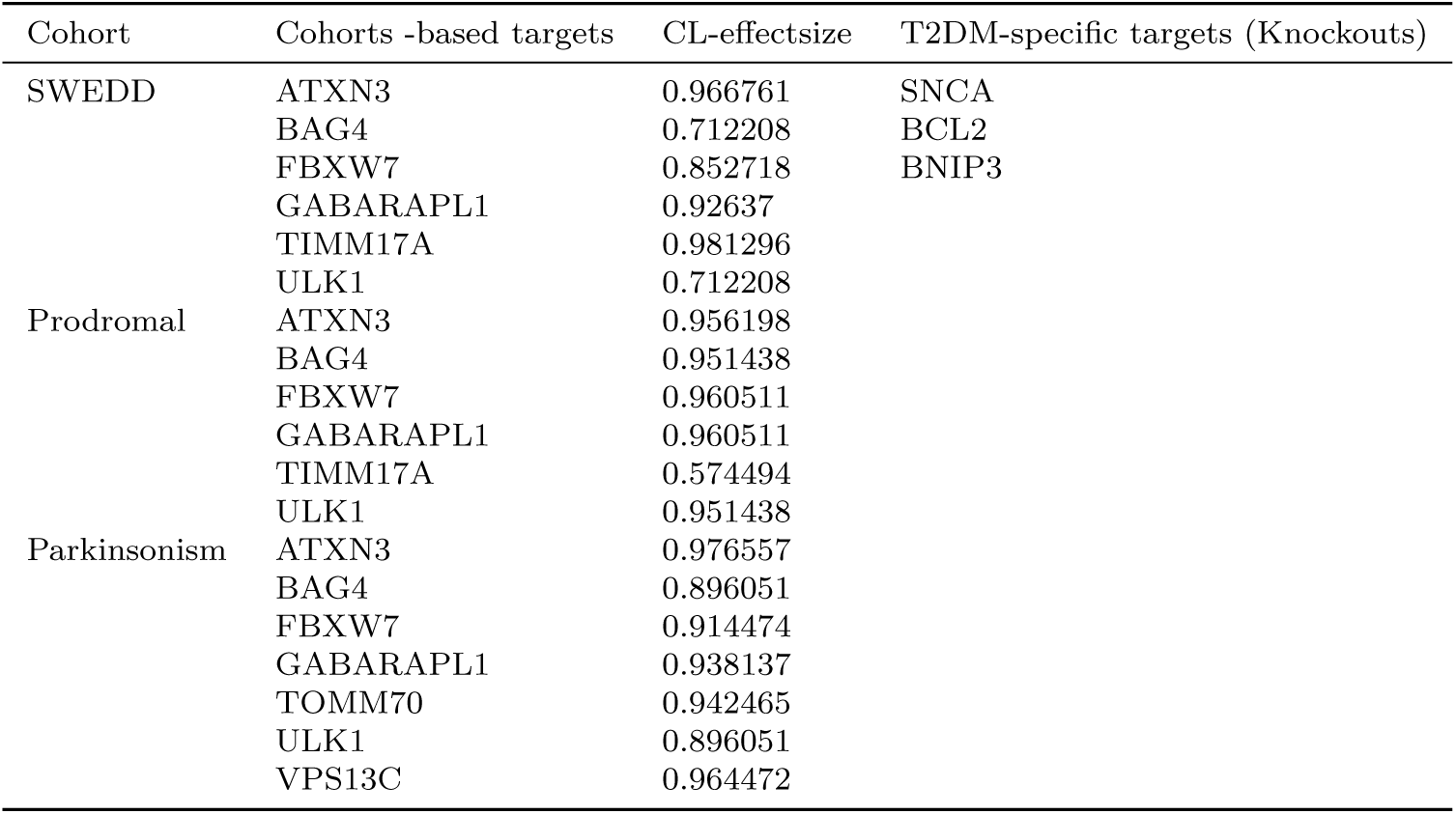
The table shows an example of parameterisation for PARKIN pathway, including two sets of parameters (miRNA based and T2DM specific which are combined) during the simulation

| Cohort | Cohorts -based targets | CL-effectsizes | T2DM-specific targets (Knockouts) |
| --- | --- | --- | --- |
| SWEDD | ATXN3 | 0.966761 | SNCA |
|  | BAG4 | 0.712208 | BCL2 |
|  | FBXW7 | 0.852718 | BNIP3 |
|  | GABARAPL1 | 0.92637 |  |
|  | TIMM17A | 0.981296 |  |
|  | ULK1 | 0.712208 |  |
| Prodromal | ATXN3 | 0.956198 |  |
|  | BAG4 | 0.951438 |  |
|  | FBXW7 | 0.960511 |  |
|  | GABARAPL1 | 0.960511 |  |
|  | TIMM17A | 0.574494 |  |
|  | ULK1 | 0.951438 |  |
| Parkinsonism | ATXN3 | 0.976557 |  |
|  | BAG4 | 0.896051 |  |
|  | FBXW7 | 0.914474 |  |
|  | GABARAPL1 | 0.938137 |  |
|  | TOMM70 | 0.942465 |  |
|  | ULK1 | 0.896051 |  |
|  | VPS13C | 0.964472 |  |

## 6 Cohort specific simulation results

Following the parameterisation, the selected BMs were analysed in two aspects:

- Investigation of molecular mechanisms in disease cohorts: parkinsonism, SWEDD and prodromal
- Effect of T2DM comorbidity in these cohorts

For each of the selected models (Table 3) two sets of results were obtained: cohort-specific and comorbidity-specific results.

### 6.1 Dopamine transcription

In ”Dopamine transcription” BM, during early simulation phases SWEDD and prodromal groups exhibited similar activation levels in mitochondrial genesis and dopamine metabolism. Conversely, the parkinsonism group manifested elevated activation levels for these biological processes in comparison to the other two groups. Noteworthy variances are also evident in neuron survival across all groups, marked by diminished activity within the SWEDD and prodromal groups (Table 7).

**Table 7.** The table presents the DTW scores in dopamine transcription BM for three different simulation stages of prodromal, SWEDD and parkinsonism. The scores are based on three different disease conditions: mitochondrial biogenesis, dopamine metabolism, and neuron survival.

| Stage | Conditions | Cohort miRNA |  |  | T2DM transcriptomics |  |  |
| --- | --- | --- | --- | --- | --- | --- | --- |
|  |  | Prodromal | SWEDD | Parkinsonism | Prodromal | SWEDD | Parkinsonism |
| Early | Mitochondrial biogenesis | 0.6771 | 0.6512 | 0.3853 | 0.3170 | 0.3185 | 0.1937 |
|  | Dopamine metabolism | 0.4592 | 0.4565 | 0.3856 | 0.1359 | 0.1322 | 0.1238 |
|  | Neuron survival | 0.3743 | 0.4894 | 0.3818 | 0.3413 | 0.3274 | 0.0668 |
| Mid | Mitochondrial biogenesis | 0.1693 | 0.1623 | 0.0493 | 0.8890 | 0.8927 | 0.0456 |
|  | Dopamine metabolism | 0.0973 | 0.0680 | 0.0496 | 0.0935 | 0.2073 | 0.0436 |
|  | Neuron survival | 0.0761 | 0.0906 | 0.0674 | 0.9226 | 0.8951 | 0.0268 |
| Late | Mitochondrial biogenesis | 0.0311 | 0.0482 | 0.0002 | 0.9676 | 0.9698 | 0.0023 |
|  | Dopamine metabolism | 0.0132 | 0.0135 | 0.0098 | 0.0364 | 0.0765 | 0.0198 |
|  | Neuron survival | 0.0110 | 0.0210 | 0.0295 | 0.0364 | 0.0765 | 0.0198 |

In the mid and late phases of simulations, we observed a general increase in activation levels for the aforementioned processes across all cohorts. An exception is the T2DM profile, where the activation levels for both mitochondrial functionality and ”neuron survival” remain constant. During these middle and late phases, distinct disparities in mitochondrial genesis arise between the SWEDD or prodromal-associated T2DM and other groups (Table 7). Within the parkinsonism-T2DM cohort, the activation intensities for ”mitochondrial biogenesis” and ”neuron survival” consistently rank lower compared to other groups across all simulation stages. These observations suggest a possible relationship between diabetes and reduced mitochondrial production and neuron survival, especially in cases of parkinsonism-T2DM.

### 6.2 Wnt-PI3K/AKT signalling

During the initial stages of simulations, a significant difference in insulin resistance emerges among the prodromal group and the SWEDD and parkinsonism groups. Both the SWEDD and parkinsonism groups exhibit similar levels of insulin resistance, in contrast to the prodromal group. The SWEDD and prodromal groups have similar levels of insulin resistance as a consequence of T2DM. The probability of the TFEB complex activation (the inactive form) is increased, whereas the active forms of TFEB (including phosphorylated TFEB and TFEB SNCA) exhibit a decreased probability of their activation levels (Table 8).

**Table 8.** The table presents the DTW scores in Wnt-PI3K/AKT BM for three different simulation stages of prodromal, SWEDD and parkinsonism. The scores are based on different disease conditions: TFEB phosphorylated, Insulin resistence, TFEB complex and Neuron death.

| Stage | Conditions | Cohort miRNA |  |  | T2DM transcriptomics |  |  |
| --- | --- | --- | --- | --- | --- | --- | --- |
|  |  | Prodromal | SWEDD | Parkinsonism | Prodromal | SWEDD | Parkinsonism |
| Early | TFEB phosphorylated | 0.9503 | 0.9215 | 0.9216 | 0.9032 | 0.9053 | 0.8998 |
|  | Insulin resistance | 0.6341 | 0.3364 | 0.3310 | 0.1111 | 0.1118 | 0.0190 |
|  | TFEB SNCA complex | 0.9622 | 0.9422 | 0.9422 | 0.8964 | 0.9215 | 0.9088 |
|  | TFEB complex | 0.3624 | 0.3572 | 0.3580 | 0.05755 | 0.0575 | 0.0125 |
|  | Neuron death | 0.3090 | 0.2714 | 0.2725 | 0 | 0 | 0.0366 |
| Mid | TFEB phosphorylated | 0.9503 | 0.9215 | 0.9216 | 0.8847 | 0.8906 | 0.8758 |
|  | Insulin resistance | 0.6341 | 0.3364 | 0.3310 | 0.0213 | 0.0203 | 0.0098 |
|  | TFEB SNCA complex | 0.9622 | 0.9422 | 0.9422 | 0.9080 | 0.9065 | 0.9078 |
|  | TFEB complex | 0.3624 | 0.3572 | 0.3580 | 0.0007 | 0.0006 | 0.0006 |
|  | Neuron death | 0.3090 | 0.2714 | 0.2725 | 0 | 0 | 0.0048 |
| Late | TFEB phosphorylated | 0.9298 | 0.9276 | 0.9243 | 0.8997 | 0.9009 | 0.8991 |
|  | Insulin resistance | 0.0241 | 0.0222 | 0.0212 | 0.0213 | 0.0203 | 0.0098 |
|  | TFEB SNCA complex | 0.9324 | 0.9279 | 0.9208 | 0.9066 | 0.8914 | 0.8931 |
|  | TFEB complex | 0.0296 | 0.0098 | 0.0097 | 0 | 0 | 0 |
|  | Neuron death | 0.0192 | 0.0005 | 0.0009 | 0 | 0 | 0 |

### 6.3 FOXO3 activity

During the early, mid and late stages of the simulation, in the prodromal group we observed increased activation of the Fission Fusion endpoint when compared to other groups. The prodromal-T2DM group had an increased activation of autophagy and oxidative stress endpoints than other groups during the three stages of the simulation. The prodromal and SWEDD groups both show a higher level of oxidative stress endpoint activation compared to the parkinsonism group. At the mid-stage of the simulation, the activation of fission and fusion within SWEDD and parkinsonism groups were similar. Furthermore, parkinsonism group displayed increased activation of autophagy relative to other groups. Also, activation of apoptosis was similar between the prodromal and SWEDD groups. In SWEDD group, activation of autophagy and apoptosis increased in the late stage when compared the prodromal and parkinsonism groups (Table 9).

**Table 9.** The table presents the DTW scores in FOXO3 BM for three different simulation stages of prodromal, SWEDD and parkinsonism. The scores are based on five different disease conditions: Response to oxidative stress, Fission Fusion, Autophagy, Mitochondrial biogenesis and Apoptosis.

| Stage | Conditions | Cohort miRNA |  |  | T2DM transcriptomics |  |  |
| --- | --- | --- | --- | --- | --- | --- | --- |
|  |  | Prodromal | SWEDD | Parkinsonism | Prodromal | SWEDD | Parkinsonism |
| Early | Response to oxidative stress | 0.6331 | 0.6220 | 0.6949 | 0.4473 | 0.4850 | 0.5954 |
|  | Fission Fusion | 0.6361 | 0.8529 | 0.8111 | 0.4676 | 0.6192 | 0.6858 |
|  | Autophagy | 0.5561 | 0.6491 | 0.5514 | 0.3789 | 0.3962 | 0.4145 |
|  | Mitochondrial biogenesis | 0.5622 | 0.5419 | 0.5824 | 0.2045 | 0.1843 | 0.1500 |
|  | Apoptosis | 0.7547 | 0.7707 | 0.5508 | 0.3947 | 0.4657 | 0.3255 |
| Mid | Response to oxidative stress | 0.5138 | 0.4952 | 0.5708 | 0.5133 | 0.4938 | 0.6417 |
|  | Fission Fusion | 0.5413 | 0.6091 | 0.6015 | 0.5188 | 0.7658 | 0.6431 |
|  | Autophagy | 0.5096 | 0.5418 | 0.4814 | 0.5195 | 0.4671 | 0.5028 |
|  | Mitochondrial biogenesis | 0.1257 | 0.1465 | 0.1633 | 0.0744 | 0.1251 | 0.0777 |
|  | Apoptosis | 0.4666 | 0.4640 | 0.3315 | 0.4231 | 0.4450 | 0.4302 |
| Late | Response to oxidative stress | 0.5099 | 0.4577 | 0.5485 | 0.5325 | 0.4703 | 0.6460 |
|  | Fission Fusion | 0.5080 | 0.5133 | 0.5469 | 0.5528 | 0.6010 | 0.6156 |
|  | Autophagy | 0.5399 | 0.4978 | 0.5367 | 0.5323 | 0.4878 | 0.5567 |
|  | Mitochondrial biogenesis | 0.0208 | 0.0372 | 0.0712 | 0.0197 | 0.0436 | 0.0392 |
|  | Apoptosis | 0.3177 | 0.2623 | 0.2752 | 0.2664 | 0.2940 | 0.3221 |

During both the mid and late stages, the SWEDD group had increased activation of oxidative stress endppoint compared to the prodromal and parkinsonism groups. The SWEDD-T2DM group also showed an increased probability of activation in oxidative stress and mitochondrial biogenesis endpoints. The prodromal group, shown increased activation of mitochondrial biogenesis endpoint. The mitochondrial biogenesis end-point within the parkinsonism group was decreased by T2DM condition compared to other cohorts (Table 9).

### 6.4 mTOR-MAPK signalling

During the early stage of the simulation, pronounced differences emerged in the activity levels of glycolysis and catabolism across all groups. Among these, in the SWEDD group we observed a change in glycolysis activation earlier and its elevated activity continued into the late stage. Conversely, the SWEDD-T2DM group showed reduced glycolysis endpoint levels at every stage of T2DM related cohorts, with an accompanying rise in catabolism and diminishing activatoin levels of glycolysis (Table 10). In the parkinsonism group, catabolic activity is more pronounced during the early and middle stages in comparison to the SWEDD and prodromal groups. Although activation of glycolysis is increased, the SWEDD group demonstrated increased levels of catabolism during the mid to late stages (Table 10). In the parkinsonism-T2DM group we observed increased activation of glycolytic activity endpoint compared to other groups with T2DM in all simulation stages (Table 10).

**Table 10.**
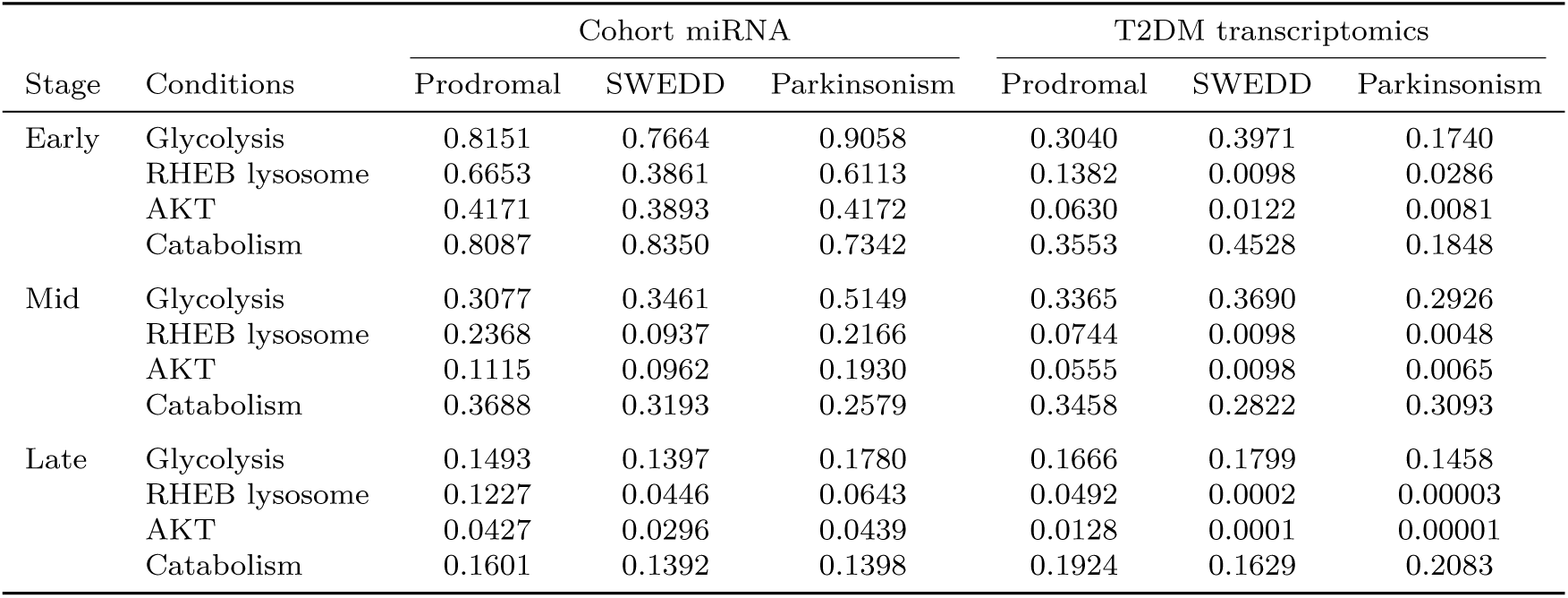
The table presents the DTW scores in mTOR BM for three different simulation stages of prodromal, SWEDD and parkinsonism. The scores are based on four different disease conditions: Glycolysis, RHEB lysosome, AKT and Autophagy.

| Stage | Conditions | Cohort miRNA |  |  | T2DM transcriptomics |  |  |
| --- | --- | --- | --- | --- | --- | --- | --- |
|  |  | Prodromal | SWEDD | Parkinsonism | Prodromal | SWEDD | Parkinsonism |
| Early | Glycolysis | 0.8151 | 0.7664 | 0.9058 | 0.3040 | 0.3971 | 0.1740 |
|  | RHEB lysosome | 0.6653 | 0.3861 | 0.6113 | 0.1382 | 0.0098 | 0.0286 |
|  | AKT | 0.4171 | 0.3893 | 0.4172 | 0.0630 | 0.0122 | 0.0081 |
|  | Catabolism | 0.8087 | 0.8350 | 0.7342 | 0.3553 | 0.4528 | 0.1848 |
| Mid | Glycolysis | 0.3077 | 0.3461 | 0.5149 | 0.3365 | 0.3690 | 0.2926 |
|  | RHEB lysosome | 0.2368 | 0.0937 | 0.2166 | 0.0744 | 0.0098 | 0.0048 |
|  | AKT | 0.1115 | 0.0962 | 0.1930 | 0.0555 | 0.0098 | 0.0065 |
|  | Catabolism | 0.3688 | 0.3193 | 0.2579 | 0.3458 | 0.2822 | 0.3093 |
| Late | Glycolysis | 0.1493 | 0.1397 | 0.1780 | 0.1666 | 0.1799 | 0.1458 |
|  | RHEB lysosome | 0.1227 | 0.0446 | 0.0643 | 0.0492 | 0.0002 | 0.00003 |
|  | AKT | 0.0427 | 0.0296 | 0.0439 | 0.0128 | 0.0001 | 0.00001 |
|  | Catabolism | 0.1601 | 0.1392 | 0.1398 | 0.1924 | 0.1629 | 0.2083 |

### 6.5 PRKN mitophagy

At all simulation stages, there was a significant difference in the activation of mitophagy across groups. Within the SWEDD and prodromal groups, mitophagy endpoint activation began earlier during the early simulation stages in comparison to the parkinsonism group. The latter group had an elevated activation of PINK1 accumulation relative to other cohorts. Additionally, the T2DM comorbidity reduced the activation of mitophagy endpoint specifically within the parkinsonism group (Table 11).

**Table 11.** The table presents the DTW scores in PRKN mitophagy BM for three different simulation stages of prodromal, SWEDD and parkinsonism. The scores are based on three different disease conditions: Mitophagy, PRKN ubiquitinated, and PINK1 accumulation.

| Stage | Conditions | Cohort miRNA |  |  | T2DM transcriptomics |  |  |
| --- | --- | --- | --- | --- | --- | --- | --- |
|  |  | Prodromal | SWEDD | Parkinsonism | Prodromal | SWEDD | Parkinsonism |
| Early | Mitophagy | 0.3303 | 0.3303 | 0.4719 | 0.0537 | 0.0537 | 0.0422 |
|  | PRKN ubiquitinated | 0.6165 | 0.6165 | 0.4545 | 0.2041 | 0.2041 | 0.0643 |
|  | PINK1 accumulation | 0.5952 | 0.5952 | 0.5870 | 0 | 0 | 0.1556 |
| Mid | Mitophagy | 0.0325 | 0.0325 | 0.0820 | 0.0078 | 0.0078 | 0.0335 |
|  | PRKN ubiquitinated | 0.1918 | 0.1918 | 0.1009 | 0.1065 | 0.1065 | 0.1212 |
|  | PINK1 accumulation | 0.1982 | 0.1982 | 0.1314 | 0 | 0 | 0.0658 |
| Late | Mitophagy | 0.0002 | 0.0002 | 0.0129 | 0 | 0 | 0.0101 |
|  | PRKN ubiquitinated | 0.0644 | 0.0644 | 0.0933 | 0.0327 | 0.0327 | 0.0534 |
|  | PINK1 accumulation | 0.0747 | 0.0747 | 0.0196 | 0 | 0 | 0.0204 |

### 6.6 Similar characteristics in the disease subgroups

Next, we compared the simulated model components among disease subgroups using the Dynamic Time Warping (DTW) algorithm (see Methods). A decreased DTW score between simulated model componets across different subgroups signifies a higher similarity in how these components unfold over time. We used the Pearson correlation coefficient to analyze DTW scores of model components across these subgroups. A Pearson correlation coefficient close to 1 indicated a strong correlation. By examining the DTW scores that are highly correlated (close to 1), we can determine a strong similarity in the temporal patterns of the analyzed subgroups. (**??**).

All results presented below demonstrate consistent patterns across the simulation stages, highlighting similarities in disease groups. The result is structured into two main points: one examining the collective trends across all disease groups, and the other focusing on pairwise comparisons between specific groups to identify the pair-wise similarities In all disease groups, we observed an increase in ”mitochondrial dysfunction” endpoint in the early stage of the disease. Further, other cellular endpoints such as ”apoptosis” and ”dopamine metabolism” are increased. As the simulation advanced to its later stages, the effect of Type 2 Diabetes Mellitus (T2DM) was observed across all groups. As a result of T2DM, the endpoints of ”neuron survival” and ”dopamine metabolism” were decreased.

As a result of comparing pairs of disease groups, similar patterns among the groups were observed as follow: i) For the prodromal and SWEDD groups, the early stages were characterized by an increase in ”mitochondrial biogenesis” endpoints influenced by T2DM and signs of insulin resistance. Additionally, We observed an increase in ”neuronal survival” and ”Autophagy” endpoints. In late stages, these groups continued to show an increase in ”mitochondrial biogenesis” endpoint associated with T2DM. Moreover, we observed an increase in ”mitophagy activity” endpoint and disruptions in ”dopamine metabolism” endpoint. ii) For the SWEDD and parkinsonism groups, the early stages demonstrated elevated levels of ”insulin resistance” endpoint. In the mid stage, we observed an increase on the ”fission metabolism” endpoint. In the late stage, an increase in ”catabolism endpoints” was observed.

## 7 Discussion

Molecular and cellular pathology in Parkinson’s disease (PD) is complex, and manifests in a range of symptoms and progression patterns, including comorbidities like Type 2 Diabetes Mellitus (T2DM) [132]. Understanding these multifaceted interactions is crucial for precise diagnostic or therapeutic approaches. For this purpose, we need systems biology approaches to investigate complex interactions between multiple molecular factors to better understand the disease mechanisms.

### 7.1 Knowledge integration

Given the variability of genetics, molecular biology, and clinical symptoms in PD, large and well-characterized cohort data is essential for identifying underlying patterns and mechanisms of the disease. For this reason, a dataset of miRNAs sequenced from the whole blood of PD patients was obtained from different disease subtypes from the PPMI study [19]. MiRNAs are considered a promising biomarker for both diagnosis and prognosis of disease because of their high stability in human fluids [20, 21]. However, they often have a broad profile of activity, so in this work we considered miRNAs having validated interactions with genes in brain tissue. We further selected only miRNAs having stable correlations with target gene expression, using curated data resources and published results related to PD, including an expression profile of PD substantia nigra [36] (see Table 1 for specific references). Most of these filtered, significant miRNAs showed downregulation in PD and were involved in mitochondrial dysfunction. Moreover, to study a potential effect of T2DM comorbidity, we analysed transcriptomic profiles of PD-related brain organoids under insulin over-exposure. Differentially Expressed Genes (DEGs) from this dataset were filtered to match the targets of cohort-specific miRNAs calculated earlier (Table 2). The common DEGs were involved in dopamine related pathways such as dopamine transcription and dopamine metabolism. This involvement suggests that insulin resistance can disrupt the normal functioning of dopamine, a key neurotransmitter in the brain. This disruption leads to impaired dopamine signaling, which is a key aspect of PD pathology. This highlights how insulin resistance contributes to the progression of PD.

Using the list of key genes compiled as above, we identified a list of pathways to construct our models. The pathways were selected from the Parkinson’s disease map (pdmap.uni.lu) [8]. The PD map is a dedicated repository of curated pathways focused on molecular pathophysiology of PD. The contents of the PD map are encoded in SBML-compliant format, allowing the construction of computational models. Thus, based on selected pathways, probabilistic BMs were created, validated for completeness and correctness, and their initial states were chosen based on the calculated miRNA effect sizes of the PPMI dataset. Boolean simulations were then performed based on these cohort-specific parameterisation. Further, T2DM data were used to parameterise the models to reflect this comorbidity. For PD-T2DM simulations, upregulated DEGs in the models were set to permanent activation, and downregulated DEGs to permanent inhibition. Such simulation allowed to separately study the effects of T2DM on PD progression.

### 7.2 Modelling-based patient stratification by disease subgroup analysis

The selected models were parameterised with cohort-based data and simulated to stratify different molecular mechanisms during disease progression. Simulations of these models represent stages in disease progression. This allowed us to compare molecular activity across the PD subtypes.

#### 7.2.1 Specific characteristics in each cohort

Prodromal cohorts exhibit molecular dysregulations that lead to higher probabilities of PD-related motor signs compared to other cohorts. These molecular dysregulations are related to impaired neuronal autophagy [133]. Further, the models show that in the prodromal cohort, mitochondrial turnover is more frequent (”Fission and Fusion” output) with higher probabilities than other cohorts in the early stages of simulation, and this pattern continues with a higher probability in the mid and late stages of simulation (Table 9). This finding is consistent with previous researches suggesting that mitochondrial abnormalities may occur early in the course of PD [134, 135]

In the SWEDD cohort, an interesting aspect is observed inhibition of ”Glycolysis and catabolism” output in the mTOR-MAPK signalling model. At the same time the protein RHEB, which has a neuroprotective role, is highly active. This suggests that RHEB may play a role in decreasing catabolic processes and potentially protecting against the development of PD [136] (Table 10). In the later stages of SWEDD, there is an increase in catabolism, which is the breakdown of molecules to release energy. This increase in catabolism is accompanied by increased glycolysis activity, which is the breakdown of glucose to produce energy. The increased catabolism and glycolysis may indicate stress adaptation. Understanding and targeting these changes could be crucial to control the PD progression.

In the parkinsonism cohort, dopamine transcription and Wnt-PI3K/AKT models show that mitochondrial biogenesis and dopamine transcription change rapidly with lower change points in the mid and late stages simulation. This finding may be due to the fact that the parkinsonism syndrome tends to progress more rapidly than other PD subgroups [137].

#### 7.2.2 Characteristics of prodromal and SWEDD cohorts

The models of SWEDD and prodromal cohorts give the similar pattern of activation in dopamine metabolism and mitochondrial biogenesis in the early stages of PD (Table 7). This finding suggests that the early stages of prodromal and SWEDD may refer to a stage at which individuals do not fulfill diagnostic criteria for clinical PD. Moreover, a recent study proposes that SWEDD patients do not have early PD [138].

The following findings suggests that even in the absence of dopaminergic neurons deficiency, as seen in SWEDD, oxidative stress may still contribute to neuronal dysfunction and warrants further investigation to fully understand its role in such conditions.[139] Dopamine transcription and Wnt-PI3K/AKT models suggest that neuronal activity endpoint may be influenced by dopamine metabolism and mitochondrial biogenesis, as these processes are important for energy production and the function of neurons. Specifically in Wnt-PI3K/AKT, SWEDD and prodromal conditions show lower levels of neuronal activity compared to parkinsonism in early stages. It is possible that dopamine metabolism is sustained in the SWEDD and prodromal early stages of simulation for longer periods of time than in the parkinsonism. As a result, dopaminergic neurons may be affected by oxidative stress, leading to a decrease in their activity [140]. Oxidative stress response may be related to dopamine metabolism and neuronal activity because oxidative stress can damage cells and disrupt normal cellular function, including dopamine metabolism and neuronal activity[140]. Prodromal and SWEDD patients exhibit similar oxidative stress responses that are higher than those observed in parkinsonism patients. This may explain the slight differences in dopamine metabolism and lower neuron survival activity in the dopamine transcription pathway observed between these two subtypes and other conditions in early stages of simulation.

Glycolysis and catabolism are central processes that are vital for the production of energy in cells. Dysregulation of these processes is observed in a wide range of disease states, including PD. Simulation results in both cohorts show that changes in glycolysis and catabolism occur earlier in SWEDD and prodromal, compared to parkinsonism (Table 10).

Mitophagy is the process of degrading and recycling mitochondria, and changes in this process may affect the function and survival of mitochondria and cells. Dysregulation of mitophagy is implicated in the development of the prodromal and SWEDD (Table 11). In the PRKN mitophagy model, the prodromal and SWEDD cohorts show higher levels of mitophagy activation than those with parkinsonism. This increase is mediated by the protein ULK1, suggesting that the process may be independent of PRKN [141], despite higher activation of ”PINK1 accumulation” in the simulation for parkinsonism cohort.

#### 7.2.3 Characteristics of T2DM comorbidity

Diabetes-parameterised and cohort-specific models demonstrate a series of differences from the results discussed above. One of the most affected is the Dopamine transcription model. It features significantly lower activation of mitochondrial biogenesis and neuronal survival at mid and late stages of simulations (Table 7). This is in line with a recent study, linking T2DM to a decline in neuron survival, mitochondrial biogenesis, and dopamine metabolism, where T2DM was associated with oxidative stress and decreased levels of dopamine and its metabolites in the striatum [132, 142]. Interestingly, in the mid and late stages, the activation of ”dopamine metabolism” endpoint is less decreased in SWEDD-T2DM cohort than the early stage of simulation (Table 7). For the Dopamine transcription model parameterised for parkinsonism-T2DM cohort, in the mid and late stages of the mitochondrial biogenesis is less decreased compared to other T2DM cohorts (Table 7).

However, in the early stages of the simulations, T2DM comorbidity is found to increase the cellular response to oxidative stress, potentially through the activation of quality control mechanisms such as Autophagy and Fission and Fusion. These processes may increase apoptosis, a form of cell death triggered in response to cellular stress [139]. Moreover, in mTOR-MAPK signalling model for parkinsonism-T2DM, we observed higher glycolysis activity and an increase in the inactivated form of RHEB and the activation of anaerobic glycolysis. This shift towards anaerobic glycolysis is thought to occur as the brain tries to maintain ion homeostasis by providing a limited amount of energy through the break-down of glucose in the absence of oxygen. However, this process ultimately leads to chemical changes that result in cell death [143–145] (Table 10). Finally, the activation of mitophagy is decreased in parkinsonism with T2DM comorbidity, and increased activation of the protein VPS13C, which delays the progression of mitophagy. In support of this, two novel cases are reported of patients who developed dementia and early onset parkinsonism in the absence of VPS13C [146].

#### 7.2.4 Common characteristics in all cohorts

The results show that dysregulation of insulin resistance is observed in all disease subgroups. Insulin resistance is a condition in which the body’s cells do not respond properly to the insulin hormone, leading to high blood sugar levels and an increased risk of diabetes and other health problems [147]. The BMs suggest that the development of insulin resistance is linked to the activity of the transcription factor TFEB (Table 8). The Boolean models show that the active forms of TFEB tend to have low activity as confirmed in[148], while the inactive form of TFEB, found in a complex with the 14-3-3 protein in the cytoplasm, tends to be elevated [149]. The 14-3-3 proteins are a family of highly expressed brain proteins with neuroprotective effects in multiple PD experimental models [149]. However, high levels of the inactive form of TFEB suggest a decrease in 14-3-3 proteins, which may increase the aggregation of alpha-synuclein and impair cellular processes, leading to insulin resistance [147]. The use of antidiabetic drugs has a beneficial role to control PD symptoms [150–152], including Metformin, suggested as a neuroprotective drug in the prodromal cohort [153]. Metformin can reduce alpha-synuclein aggregation and improve cellular processes associated with age-related conditions [17, 152]. The Boolean model suggest that dysregulation of TFEB and its regulated genes plays an important role in insulin resistance and controlling mitochondrial function in PD. A recent study shows that abnormalities in TFEB cause a failure of endolysosomal and autophagic pathways[154].

Our BMs can help to explain hypothesis to understand complex diseases, and propose better therapies and diagnostics. Boolean modelling approach allows comparing the model attractors to the disease signature and design perturbation experiments that cause transition of the pathological signature towards a healthy state. As discussed, the results and hypothesis generated by the models were inline with the existing literature findings. The models proposed that dysregulation of cellular phenotypes (model endpoints) varies among different disease cohorts. The results of the Boolean models can be used to improve similarity-based differential diagnosis in PD. This can be achieved by identifying the common cross-talk between different subtypes of the disease. PD has different subtypes that may be presented with similar phenotypes (endpoints), but have different underlying causes/regulators. As a result, more precise therapeutic strategies need to be developed based on different causes even if they share the same symptoms. This explains that the targets and treatment strategies should be tailored to each disease subgroup. It could be possible to construct experimental models representing each subgroup, and perturb the targets predicted by the model to observe the pathological signatures.

### 7.3 Limitations of the study

Our work faces a number of limitations. First, the stratification of the models relies on miRNAs and in this work, the specificity of miRNAs is validated in independent studies. Indeed, some miRNAs were found to have mismatched expression levels between the PPMI and literatue (Table 1), which may stem from various factors such as the presence of other miRNAs, the availability of specific transcription factors, and the overall gene expression profile of the cell [22]. Also, to simulate the mutation effect of T2DM on PD cohorts, we use a snapshot of dynamic data which is a limited representation of a complex comorbidity of PD. A more comprehensive approach would require analysis of data describing molecular profiles of progression in both disorders over time.

Next, a number of pathway-based models are analysed separately, while in fact they are interconnected. The integration of pathways may allow better understand the disease progression and therapeutic responses of PD, which requires a broader investigation is necessary.

Another limitation is the granularity of model parameterisation, as we only focused on disease subtypes, without considering other factors that may affect the dynamics of the disease, such as gender and age. Finally, our work lacks experimental validation of proposed combinatorial interventions, limiting to supporting literature findings.

## 8 Conclusion

Probabilistic Boolean Models (PBMs) are promising tools to advance research hypotheses by simulating complex pathways based on available data. These models are effective in capturing the stochastic nature of molecular interactions, and offer insight into their dynamics. In our work, PBMs integrated with empirical data helped to identify critical variation in pathways across PD subtypes, and suggesting the development of targeted therapies. However, our PBMs are based on assumptions and simplifications of complex biological processes involved in PD. Although this makes them more tractable and easier to analyze, they may not accurately reflect the complexity of the relationships between different factors, especially when data is incomplete. To improve the use of PBMs in clinical applications, it is important to focus on improving data integration from various sources, such as expression data, interaction data, and literature-based knowledge. This will require the development of robust and scalable methods for data integration, as well as the establishment of standards for data representation and interoperability.

## Declarations

- Funding This work was supported by the funding from the European Union’s Horizon 2020 research and innovation programme under grant agreement No. 733100: SYSCID—A systems medicine approach to chronic inflammatory diseases.
- Conflict of interest/Competing interests The authors declare that the research was conducted in the absence of any commercial or financial relationships that could be construed as a potential conflict of interest.
- Authors’ contributions AH: investigation, conceptualization, writing–original draft. VS: supervision, writing–review and editing. RS: supervision, writing–review and editing. MO: conceptualization, review and editing, supervision. All authors contributed to the article and approved the submitted version.

